# Arousal state fluctuations are a source of internal noise underlying age-related declines in speech intelligibility

**DOI:** 10.1101/2025.05.09.653191

**Authors:** Samuel S. Smith, Jenna A. Sugai, Daniel B. Polley

**Author notes:** Correspondence: Samuel S. Smith, **Email:**. **Competing Interest Statement:** The authors disclose no competing interests.

## Abstract

Understanding speech in noisy, multi-talker environments is crucial for social communication but becomes increasingly challenging and frustrating as we age. Here, we simulated the acoustic challenges of multi-talker listening and found that adults over 50 years old (N = 76) recognized speech more slowly, less accurately, and less consistently than younger adults (N = 107). While peripheral hearing status accounted for average differences in speech intelligibility by age, it did not account for moment-to-moment variability in speed and accuracy - fluctuations central to the frustration experienced by older listeners in challenging environments. We hypothesized that age-related changes in brain arousal systems might account for the fluctuant “noise” in speech processing observed in older listeners. To isolate the contribution of arousal state independent of hearing status and cognitive load, we measured the pre-stimulus pupil-indexed arousal state (PPAS) immediately prior to speech onset. Older – but not younger - adults exhibited a striking inverted-U relationship between PPAS and speech recognition accuracy. Notably, pupil-indexed listening effort measured seconds later during speech encoding was not associated with trial-to-trial performance. Moreover, older adults exhibited altered arousal regulation, occupying a lower PPAS extremum not observed in younger listeners that was specifically associated with performance deficits and subjective listening difficulties reported in hearing health questionnaires.

These findings show that age-related changes in central arousal states interact with peripheral hearing status to offer a more complete explanation for why older adults find speech processing in social setting so challenging.

**Significance Statement:** As we age, following a conversation in crowded environments becomes more difficult and frustrating, even when hearing tests appear normal. This study shows that variations in arousal level (measured through pre-stimulus pupil size) are tightly linked to fluctuations in the speed and accuracy of speech processing in adults over 50 years of age. Changes in arousal state regulation uniquely accounted for the moment-to-moment variations in speech intelligibility and were more closely associated with self-reported listening challenges than other conventional measures. These findings highlight the contribution of brain-wide arousal systems to real-world listening challenges and identify a straightforward measure that could be used to more comprehensively assess and potentially improve speech understanding in social settings.

## Introduction

The auditory system supports effortless and near-instantaneous comprehension of a conversational partner’s speech. This routine act is made all the more remarkable when, as a result of aging, speech recognition accuracy declines, and greater levels of effortful listening are required. Age-related declines in speech intelligibility are particularly common in reverberant interior spaces with multiple competing speakers (1, 2), as would be encountered in a typical office conference room or busy restaurant. Age-related difficulties with speech understanding in social multi-talker settings may incur broader neurological (3, 4) and societal (5, 6) costs, highlighting the need for a more complete understanding of the underlying causes and the development of assistive listening devices that better address the challenges of speech intelligibility in crowded multi-talker settings (7–9).

Age-related declines in multi-talker speech intelligibility have been associated with a cascade of degenerative and compensatory changes in the auditory periphery, central auditory encoding, and auditory cognition. Compromised middle ear mechanics and degeneration of sensory and non-sensory cells in the cochlea are major contributors to elevated hearing thresholds (10, 11). Primary afferent neurons also exhibit precocious degeneration, effectively breaking the essential synaptic connection between inner hair cells and the auditory nerve years before age-related loss of hair cells and elevated hearing thresholds are noted (12–14). Selective cochlear neural degeneration has also been associated with impaired hearing in noise (15, 16). Peripheral afferent degeneration tips the balance of excitation and inhibition in the central auditory pathway towards disinhibition and hyperexcitability, which can interfere with high-fidelity temporal processing and perceptual object segregation (17–20). At the level of auditory cognition, aging and hearing loss compromise the deployment of cognitive resources required in complex listening environments, such as attentional control, working memory, and effort (21–26).

Whilst current models identify diverse causes of speech intelligibility errors, they are all predicated on the assumption that listening errors arise from peripheral and central processing of the sound stimulus itself. Yet, from one moment to the next, internal brain states that host bottom-up sensory representations are subject to stochastic fluctuations regulated by diffuse neuromodulatory networks (27– 29). Though trial-to-trial variability is typically treated as experimental noise and averaged across, it may capture important underlying perceptual processes such as top-down cognitive influences. The Yerkes-Dodson law asserts that perceptual errors are more frequent at lower and higher levels of arousal (30), loosely associated with drowsiness and anxiety in the extremes. In the rodent auditory cortex, these extremes have been correlated with slow neural oscillations and depolarized membrane potentials in layer 4/5 auditory cortical neurons respectively (31). The central neuromodulatory systems that modulate the membrane potential of cortical neurons and behavioral performance during challenging listening tasks are also the upstream regulators of the ciliary ganglion that control the pupil dilatory musculature (32, 33). In this way, the pupil mirrors internal states of arousal and exhibits a non-monotonic “inverted-U” relationship between arousal and performance as described by the Yerkes-Dodson law (i.e., optimal performance at intermediate, rather than small and large pupil diameters) (31, 34–38).

Together, these findings motivate and inform the central thesis of our study: that fluctuations in pupil-indexed arousal state can – independent of any speech encoding process – predict the trial-to-trial variations in speech recognition outcomes, particularly in older adults for whom listening errors are common. To avoid our interpretation being contaminated by stimulus-related and motor responses, we quantified pupil sizes in the short windows of time just prior to the delivery of multi-talker stimuli. To minimize the influence of the test itself on arousal levels and fatigue (39), we limited ourselves to a minimal test battery (15-20 minutes) performed on a relatively large sample (183 study participants).

## Results

### Performance is impaired at low and high levels of pupil-linked arousal in older adults

All participants gave informed consent for our study and completed an eight-block closed-set speech recognition task alongside measurements of pupil diameter. To distill the challenge of listening in multi-talker settings, participants were asked to report four digits vocalized by a target speaker (40). The four digits were simultaneously masked by two additional speakers at a relatively easy (9 dB) or difficult (0 dB) signal-to-noise ratio (SNR) (**Fig. 1A**). While study participants were close to ceiling during 9 dB SNR trials, we observed that performance at 0 dB SNR was lower and notably variable across trials (**Fig. 1B**). Trial-to-trial variability was more dispersed than anticipated by a stationary binomial process (one-sample t-test of dispersion factors, *P*<0.001, N=183), pointing to fluctuating influences on speech intelligibility that cannot be observed with a grand average. Here, we examined whether spontaneous shifts in arousal corresponded with variations in multi-talker speech intelligibility, measuring the pre-stimulus pupil-indexed arousal state (PPAS) during a 1s period before the first digit of each multi-talker stimulus (**Fig. 1C**).

**Figure 1.**
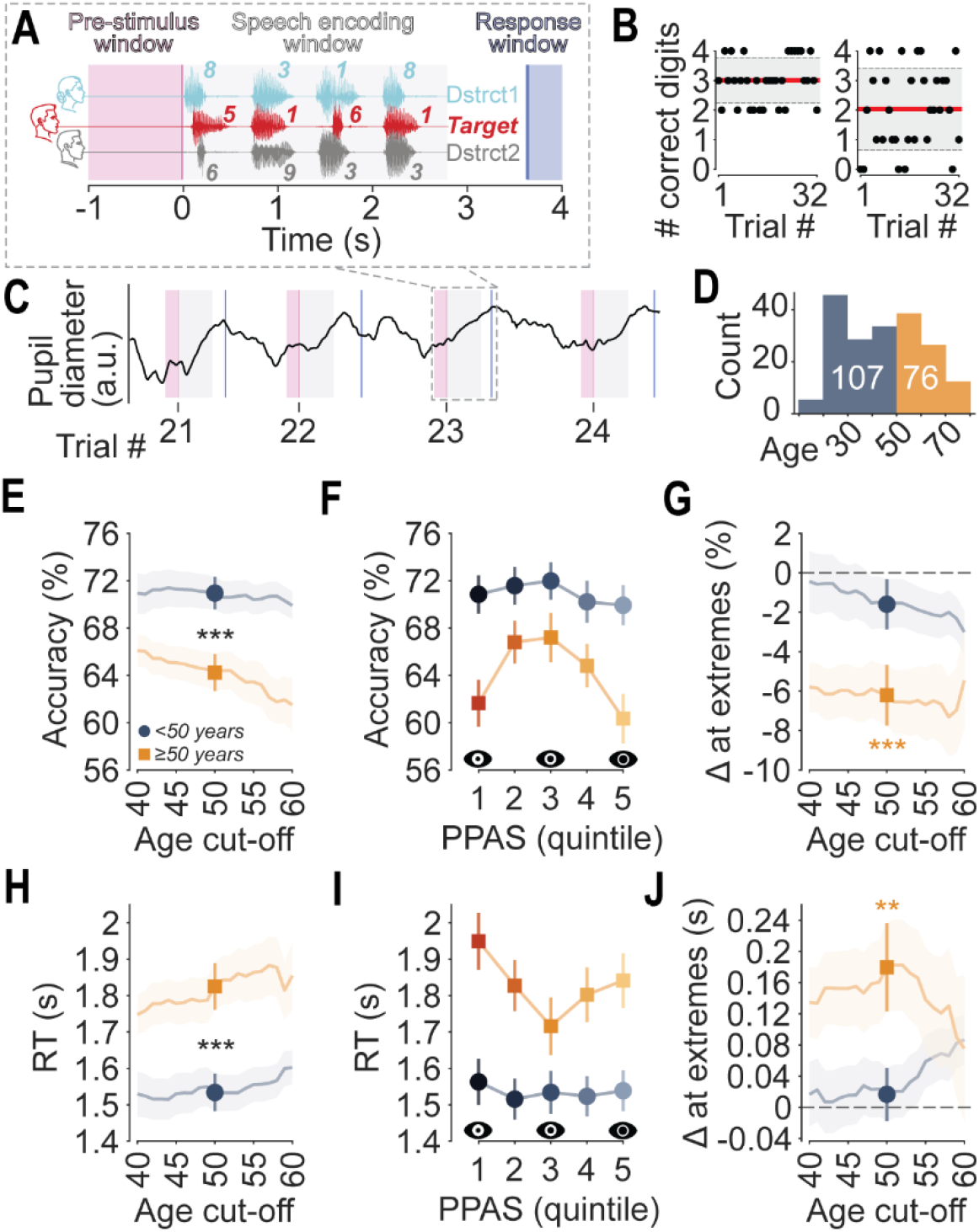
Performance is impaired at low and high levels of pupil-linked arousal in older adults. (A) Schematic of the multi-talker digit recognition task in which study participants were asked to report four digits vocalized by a target speaker simultaneously masked by two additional distracting speakers (Dstrct1, Dstrct2). (B) Outcomes on individual 0 dB SNR trials for two study participants, highlighting the presence of trial-to-trial variance. The red line shows mean performance and the grey shaded region the standard deviation. (C) An example pupil trace recorded during a 0 dB SNR block. Trial numbers are shown along the abscissa and pre-stimulus window are underlaid in pink. (D) A histogram of study participants split at age 50 into younger and older adult cohorts. (E) Older adults (orange square) correctly reported a significantly smaller proportion of digits in comparison to younger adults (blue circle) during 0 dB SNR trials (two-sample t-test, *P*<0.001, N=107/76). This trend generalized for age cut-offs above and below 50. Error bars/shaded region denote standard error. (F) Dependence of task accuracy on pre-stimulus pupil-indexed arousal state (PPAS) during 0 dB SNR trials. To normalize for variations in pupil size across individuals and age, PPAS values were assigned to one of five equally sized bins per person. Smaller and larger diameters are associated with increased errors in older adults. (G) We measured differences in accuracy between the 3^rd^ and 1^st^/5^th^ quintiles (Δ at extremes). Accuracy at pupil-indexed extremes significantly decreased in the older cohort (one-sample t-test, *P*<0.001, N=76), and were again robust to the exact choice of age cut-off. (H-J) as (E-G) for response times (RT). Older adults took significantly longer to respond than younger adults (two-sample t-test, *P*<0.001, N=107/76) with increased response times at pupil-indexed extremes during 0 dB SNR trials (one-sample t-test, *P*=0.002, N=76).

As expected, we found that when averaging across the more difficult 0 dB SNR trials older adults performed significantly worse (**Fig. 1D,E**; statistical reporting provided in figure legend). Strikingly, for the cohort of older adults, we observed an inverted-U relationship between PPAS and task performance, such that speech recognition errors were more likely during both smaller and larger pre-stimulus pupil sizes (**Fig. 1F**). We quantified this as the difference in performance between the central arousal state (i.e., the 3^rd^ quintile) and the extremes (i.e., the 1^st^ and 5^th^ quintiles) (**Fig. 1G**). We also found that response times in older adults were longer at lower/higher levels of PPAS (**Fig. 1H-J**). PPAS-related reductions in accuracy were correlated with PPAS-related increases in response times (Pearson’s r=-0.31, p<0.001, N=183). Interestingly, PPAS was unrelated to listening errors in younger adults.

As expected, hearing thresholds were elevated in our cohort of older adults, and the degree of hearing loss mediated the overall influence of age on speech intelligibility (**Fig. S1A-C**). However, the degree of hearing loss could not mediate the increased effect of pre-trial arousal state on speech intelligibility (**Fig. S1D**). We also fit a generalized linear mixed-effects model to trial-level binomial outcomes. Like in the mediation analysis, we saw that PTA outstripped age as a predictor of poorer speech intelligibility (**Fig. S2, Table S1**). But the influence of pre-trial arousal state on speech intelligibility only interacted with age. These findings indicate that age-related changes in how arousal influences speech intelligibility appear to be separable from the influence of age-related presbycusis.

### Errors are due to reporting competing speech

Not all speech errors are alike. For example, the listener may perceive a word that was not uttered or may misattribute the speech from a competing background speaker to the speech stream of the target speaker. We found that masker confusion – i.e., confusing the speech of the distracting speaker with the target speaker – accounted for most of the performance differences between younger and older adults, as well as the inverted-U relationship with arousal (**Fig. S3A**). This result indicated that the digit stimuli at lower/higher levels of pupil-linked arousal were accurately encoded but improperly classified. We also found that PPAS measured just prior to the first digit was associated as closely with performance on the first digit as the fourth digit, suggesting a prolonged influence of arousal state over multiple seconds (**Fig. S3B**).

### Older adults exhibit a broader range of pupil-indexed arousal

How does the influence of arousal on speech intelligibility change with age? A parsimonious explanation is that the dynamics and stability of arousal states themselves wane with age. For example, pupil size is known to reduce with time-on-task related fatigue, and this could become accentuated with age (41–43). Although our task was relatively short (15-20 minutes), in younger adults the smallest PPAS quintile was 12% more likely by the end of the task. However, this trend was not accentuated in older adults and we in fact observed no significant difference between the start and end of the task (**Fig. 2A,B**). Given that only older adults showed a relationship between PPAS and task performance, this seems unlikely to be due to pupil-indexed fatigue. We also considered whether older adults might exhibit more volatile arousal states. However, state transition statistics did not differ from younger adults (**Fig. 2C**).

**Figure 2.**
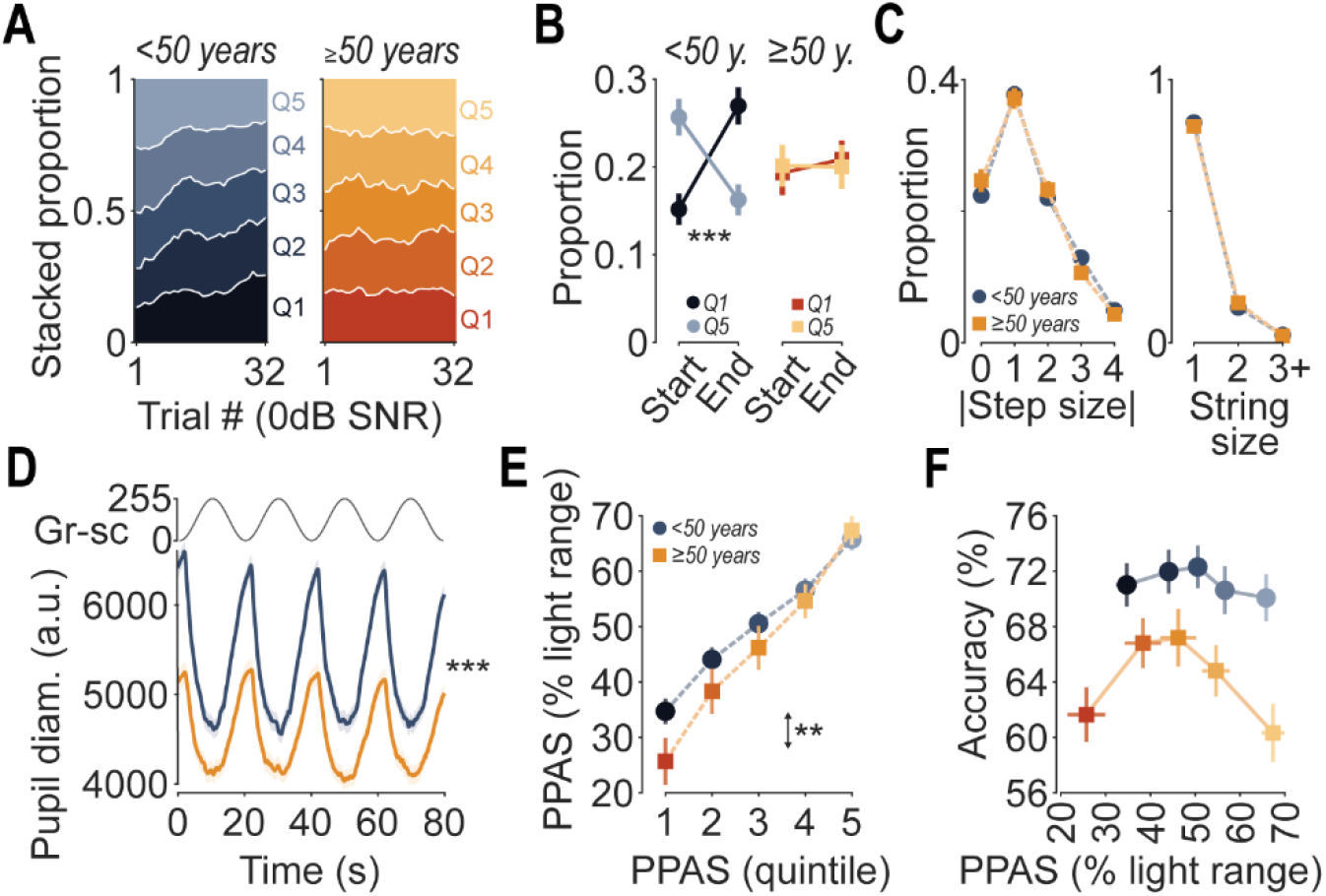
Older adults exhibit a broader range of pupil-indexed arousal. (A) Stacked plots show pupil quintile likelihoods as a variable of trial number (8 trial moving average). Younger and older adults are shown in blue and orange, respectively, with smaller pupil sizes indicated by quintiles of a darker shade. (B) The proportion of trials spent in the smallest pupil quintile (Q1) increased between the start and the end of the experiment for younger adults (paired t-test, *P*<0.001, N=106), but not older (paired t-test, *P*=0.68, N=73). (C) State transition statistics for younger versus older adults. Proportions are shown for the magnitude of step sizes between consecutive trials (left) alongside the number of consecutive trials spent in given state (right). (D) Pupil diameter (Pupil diam.) was measured during an alternating grayscale (Gr-sc) light stimulus, from which ocular limits of dilation and constriction were estimated for each participant. Consistent with reported trends, the ocular limits of the pupil in response to light was significantly smaller in older adults than younger (two-sample t-test, *P*<0.001, N=102/72; grayscale). (E) PPAS, as measured relative to light responses, were broader and lower in older adults (two-sided Wilcoxon rank sum test, *P*=0.005, N=102/72). (F) Dependence of behavioral performance on PPAS during 0 dB SNR trials, as measured relative to the light response bounds. Error bars denote standard error.

Another explanation is that older adults might experience different extremities of arousal. To assess this, we benchmarked pupil-linked arousal against the ocular limits of each participant’s pupillary light reflex. In this way we control for individual variations in pupil size, as well as age-related miosis, in which the pupil is known to be smaller in older adults (**Fig. 2D**; (44). We then referenced each participant’s pre-stimulus pupil sizes to their maximally constricted and dilated sizes, as recorded during the light stimulus (denoted as 0% and 100%, respectively). We found that older adults sampled a broader range of pupil-linked arousal levels, skewed towards smaller sizes (**Fig. 2E**). Re-plotted task performance on PPAS benchmarked against light responses revealed that younger adults did not exhibit the lower extents of pupil-linked arousal where speech intelligibility errors were especially common in older adults (**Fig. 2F**). Again, this is unlikely a result of task-related fatigue because in older adults the lower extents of pupil-linked arousal were as common at the start of the experiment as at the end (Fig. 2A,B, Fig S3A; more so, performance accuracy was higher at the end of the task than at the start [Fig S3B]).

### Pupil-indexed listening effort is greater in older adults but does not account for trial-to-trial variation

Arousal is one of several cognitive processes thought to shape speech perception. Amongst these, larger pupil dilations during the active parsing of speech have been shown to reflect increased levels of listening effort - defined as the allocation of mental resources in the pursuit of a listening-related goal (23, 24, 40). Here, we confirmed that stimulus-evoked pupil responses were larger in more difficult listening conditions, i.e., 0 dB SNR versus 9 dB SNR (**Fig. 3A**). In accordance with previous work, we also observed that evoked pupil responses were larger in older adults relative to younger adults (45, 46).

**Figure 3.**
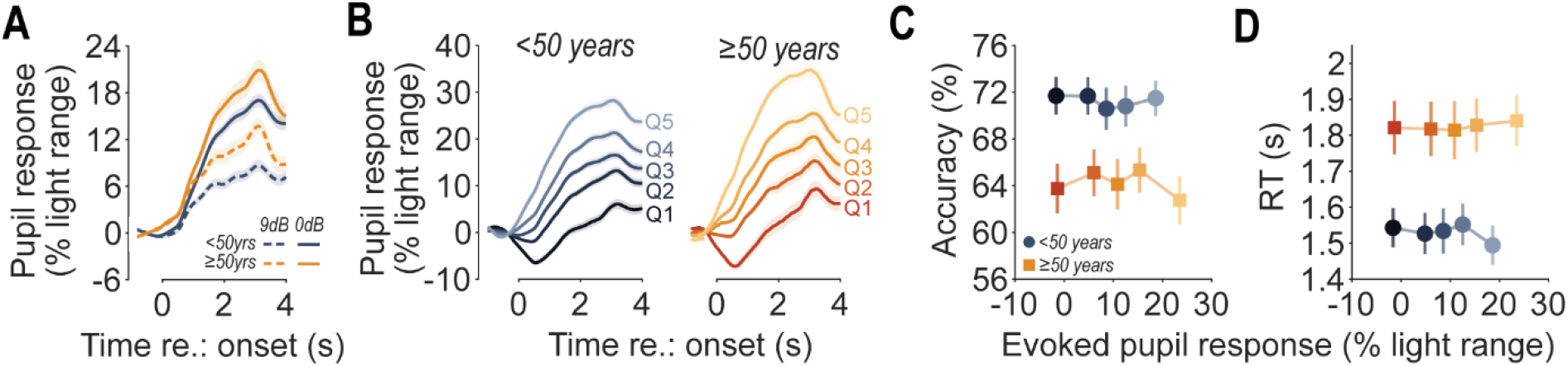
Pupil-indexed listening effort is greater in older adults but does not account for trial-to-trial variations. (A) Stimulus-evoked pupil responses were larger in more difficult listening conditions and in older adults (two-way ANOVA, N=102/72, main effect of SNR [F= 207.1, *P*<0.001], main effect of age [F=10.8, *P*=0.001], SNR x age [F=1.0, *P*=0.32]). (B) In the same way that we partitioned pre-stimulus pupil diameters into quintiles, stimulus-evoked pupil responses were assigned to one of five equally sized bins per person. (C, D) No relationship was observed between accuracy or response times (RT) with stimulus-evoked pupil responses.

Pupil-indexed listening effort is generally measured as the average over a block of trials. If pupil-indexed listening effort shaped speech outcomes as widely assumed, then trial-to-trial variations in pupil-indexed listening effort should account for trial-to-trial fluctuations in multi-talker speech recognition performance. For example, rises and falls in listening effort could influence active speech inference mechanisms, which could be tied to or independent of arousal levels. Repeating the same type of analysis that we previously applied to the pre-stimulus pupil data, we assessed the dependence of accuracy and response times on the size of the stimulus-evoked pupil response (i.e., changes relative to the pre-stimulus pupil size) (**Fig. 3B**). Neither digit recognition accuracy nor response times were found to correspond with stimulus-evoked pupil size (**Fig. 3C,D**). This indicates that pupil-linked listening effort is not determinative of within-person fluctuations in listening performance but more a product of stimulus characteristics and demographic qualities such as age.

### Deficits at low arousal levels correlate with self-reporting listening experience

The digits task assesses psychoacoustic aptitude in active listening environments with multiple speakers. However, the extent to which this relates to the lived experience of communication issues in everyday life is less well established. To this end, all participants completed the Speech, Spatial and Qualities of Hearing Scale (SSQ; (47)) a 49-item questionnaire designed to assess hearing handicap across multiple domains. Task performance and SSQ scores had a correlation coefficient of 0.18 (*P*=0.017, N=179) (**Fig. 4A**). This correlation could speak to the subjectivity of self-report questionnaires, but it might also indicate that the experience of hearing difficulties is not fully explained by bottom-up acoustic features like signal-to-noise ratio. Instead, challenges aligning optimal arousal states with listening demands could be as – or more – related to the hearing challenges referenced in questionnaires like the SSQ.

**Figure 4.**
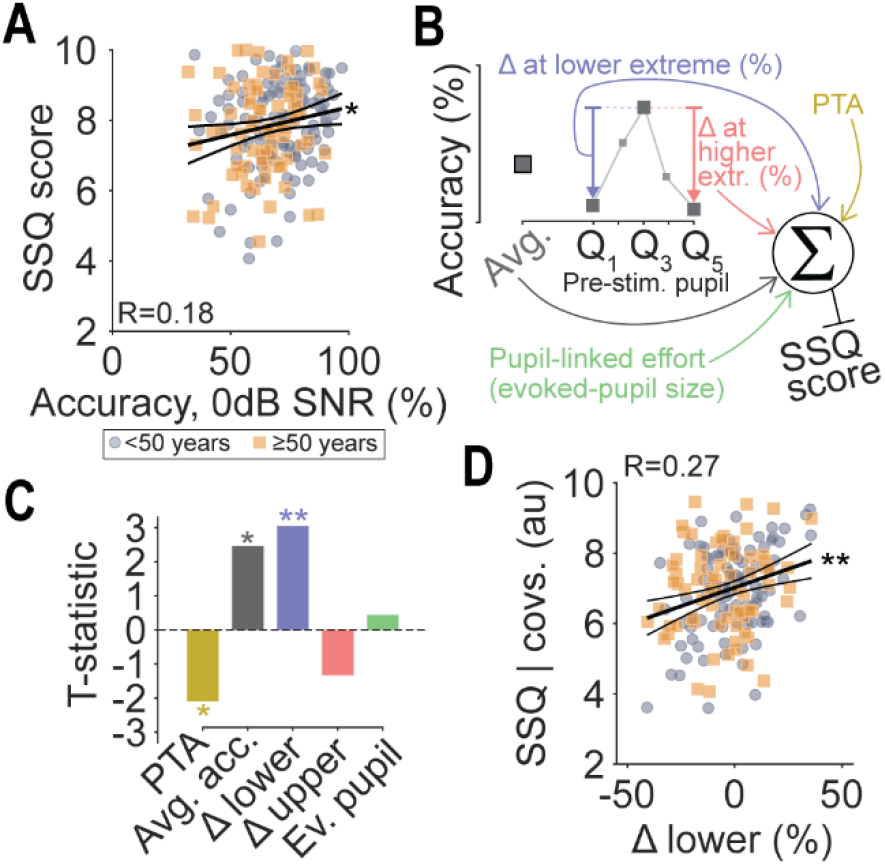
Deficits at low arousal levels correlate with self-reported listening experience. (A) Average digits-in-noise performance at 0 dB SNR and Speech, Spatial and Qualities of Hearing Scale (SSQ) scores (Pearson’s r=0.18, *P*=0.017, N=179). Blue circles denote adults younger than 50 years and orange squares denote adults 50 years or older. (B) A schematic of the regression model for SSQ scores. (C) T-statistics for each coefficient in the model predicting SSQ scores (pure-tone average [PTA], mean accuracy [Avg. acc.], mean change in accuracy between the 3^rd^ and 1^st^ pre-stimulus pupil quintiles [Δ lower], mean change in accuracy between the 3^rd^ and 5^th^ pre-stimulus pupil quintiles [Δ upper], mean evoked pupil response [Ev. pupil]). (D) Deficits at lower levels of pupil-linked arousal correlated with SSQ scores, shown after controlling for other model co-variates (covs.; Pearson’s r=0.27, *P*=0.002, N=170).

To investigate this, we built a regression model to predict individual SSQ scores (**Fig. 4B**). The model incorporated average performance on the task as an explanatory variable and, importantly, individual changes in performance at the extremes of PPAS (i.e., performance decreases/increases for trials in which the pre-stimulus pupil size belonged to the smallest or largest quintiles, respectively). We also included pure-tone averages and pupil-linked effort as potential explanatory factors. We found that the dominant predictor of poorer hearing outcomes on the SSQ was whether individuals exhibited large performance deficits in the lower extremes of pupil-indexed arousal (**Fig. 4C**,**D**). This predictor was more influential than any other in the model, and approximately 1.4x more influential than PTA (based on standardized model coefficients).

## Discussion

Here, we investigated the role of arousal during a psychophysical multi-talker speech recognition task. By focusing our analysis on pupil size measured prior to speech onset, we could isolate arousal from other pupil-associated cognitive operations leveraged during and following parsing of speech, such as listening effort and memory (23, 31, 48, 49). We found that, in older adults – but not younger adults - multi-talker intelligibility depended on PPAS, underlining a prominent influence of arousal state in this population. This finding identifies a cognitive signature, pupil-linked arousal, that is associated with age-related decline and contributes to communication difficulties.

### The neural and cognitive actions of pupil-linked arousal

Pupil diameter is a one-dimensional signal that is indirectly regulated by neuromodulatory centers throughout the brain, including locus coeruleus noradrenergic neurons (50–53), cholinergic neurons of the basal forebrain (50, 54, 55), orexinergic neurons of the hypothalamus (32, 56), and serotonergic neurons in the Dorsal Raphe (57). Given the heterogeneity of upstream regulators, it comes as no surprise that pupil diameter has been associated with an equally wide range of cognitive processes including listening effort (23, 24, 40), affective processing (58), working memory (59, 60), imagination (61), selective attention (35, 62), expectation violations (63, 64), and individual biases (65). Importantly, we found that pupil diameter measured one second before speech onset strongly predicted task performance while measuring the pupil just seconds later, during speech processing had no association with task performance. One interpretation of this finding is that pre-stimulus pupil measurements provide a relatively direct measure of arousal while post-stimulus pupil reflect a more heterogenous combination of influences related to arousal, executive control, and memory processes. The temporal specificity of pupil-based predictions of single trial task performance argues against the characterization of the pupil as a catchall measure and instead underscores the value of leveraging time within an experimental trial to infer the relationship between pupil size and the cognitive and neuromodulatory states that regulate task performance.

We found that PPAS clearly had an inverted-U relationship with multi-talker speech intelligibility, such that errors most commonly occurred during the most and least dilated states. Generally, the Yerkes-Dodson law (30) has been employed to explain these dynamics, and while the inverted-U relationship between pupil-indexed arousal and speech intelligibility has not been described (42, 43), the non-monotonic association between pupil diameter and task performance has been reported in a wide range of tasks and species (31, 34–38). In our study we saw that most errors produced at the pupil-indexed extremes of arousal were due to misreporting of digit stimuli spoken by competing speakers. This would suggest that pupil-linked arousal may reflect interference with stream formation and attentional processes (21, 22, 66, 67), as opposed to stimulus encoding fidelity and perceptual salience.

Task design can significantly impact the contribution of arousal states to measurements of brain and behavior. One motivation for our study was to isolate more rapid fluctuations in arousal states from the slower, monotonic changes in arousal that come with extended testing on rather dull laboratory-based psychophysical tasks. We hoped to accomplish this by recruiting a sample size that was 5-10 times larger than most studies but restrict the testing to approximately 15-20 minutes, thereby imitating the listening challenges in realistic conversation durations with minimal contributions of slower, global changes in vigilance. Prior studies employed longer experiments with hundreds or thousands of trials, which likely would have changed the range of distribution of arousal states during testing. Indeed, we found that younger adults in our study exhibited a narrower range of pupil-indexed arousal states than the older adults and did not exhibit the same inverted-U relationship with accuracy. Other studies have also reported deviations from the inverted-U, such as linear relationships that may be explained by under-sampling the Yerkes-Dodson curve through constraints related to task design, such as task difficulty, modality, and choice of behavioral measure (37, 43, 68, 69).

### Clinical significance

Sensory decline is becoming more prevalent in an aging population, with notable societal and economic implications (3, 4, 70). For one of the most reported issues – hearing difficulties in noisy environments – current diagnostic tests are insufficient. Even efforts to incorporate behavioral speech-in-noise testing might fall short. Older adults who report struggling in busy and loud settings may still have good scores on speech-in-noise tests, by utilizing top-down resources to compensate for reduced peripheral encoding (25). Sufficient speech understanding may occur in spite of increased levels of mental burden and tiredness. For example, perception of acoustically degraded signals has been shown to draw on resources otherwise reserved for memory and comprehension (24, 59). That our measure capturing arousal-related deficits correlated better with SSQ scores than average task performance speaks to this. However, our summary measure was a relatively ad-hoc one, and tests that deliver more precise and fuller characterization of pupil-arousal profiles may better serve as assays for these otherwise intangible qualities of listening.

## Materials and Methods

### Participants

Data presented are from 183 adults between 18 and 76 (mean age = 44.4 years, 103 female) that were fluent in English and provided their written informed consent to participate in the study. Participants were recruited through flyers, word of mouth, clinician referrals from the Mass Eye and Ear otolaryngology clinic, and by posting to the Mass General Brigham participant recruitment website. All procedures were approved by the Mass General Brigham Institutional Review Board (#IRB2019P002423) and took place at Mass Eye and Ear. Participants underwent extensive screening for medical history, cognitive function, and mental health. Participants also underwent audiological evaluation. Air conduction thresholds were measured for each ear with insert earphones (EarTone-3A, Oaktree Products, Chesterfield MO) for pure tones ranging from 0.25 – 8 kHz in octave intervals using the modified Hughson-Westlake procedure. Thirty-two participants were excluded based on the screening process (ten did not pass mental and cognitive screeners, twenty-two did not pass audiometric testing). Two participants were excluded due to confounding ophthalmology-related medical issues. Twenty-eight participants did not successfully complete the multi-talker digits testing (fifteen dropped out following screening, thirteen did not complete all blocks). Of the remaining 205 participants that successfully completed the multi-talker digits testing, eighteen had data excluded due to inordinately high rates of pre-stimulus blinking (see “Pupillometry data analysis”) and four due to unusually low task performance at the easier signal-to-noise ratio (less than 90% accuracy at 9 dB SNR).

### Acoustic stimulus generation

Acoustic stimuli were generated at 48828 Hz in LabVIEW (National Instruments, Austin TX) with a 24-bit D/A converter (TDT RX6, Tucker-Davis Technologies, Alachua FL), passed through a unity-gain power amplifier (TDT SA1). Earphones were calibrated following American National Standards Institute (ANSI) specifications. Insert earphones were connected to a 2-cc acoustic coupler with rigid tube attachment (AEC202, PCB Electronics, Depew NY) fitted with a ½” microphone (Larson Davis Model 2541, PCB Electronics). The microphone preamplifier (Larson Davis Model 2200) was connected to a 24-bit A/D input on the TDT RX6. Custom LabVIEW software measured sound pressure level (in dB SPL) as a function of frequency in response to 20-ms chirp stimuli (500 averages).

### Pupillary light reflex

Participants’ heads were stabilized throughout the session with a padded head support frame with an adjustable chin and forehead rest (SR Research Ltd., Ottawa CA). A 20 second light stimulus sinusoidally alternating between maximally bright and dark screens was presented for four consecutive repeats. Participants fixated towards the center of the front-facing monitor whilst pupil diameter was simultaneously recorded with the EyeLink 1000 Plus (SR Research Ltd.) at a sampling rate of 1 kHz.

### Multi-talker digits task

All testing was performed in a sound attenuating RF-shielded booth measuring 2.9m x 2m x 2m (L x W x H; Eckel Industries Inc., Ayer, MA) with an ambient noise level less than 16.4 dBA. In a given trial, participants were required to report four digits that were spoken by a male target speaker (F_0_=115 Hz) masked by two additional speakers (F_0_=90 Hz, F_0_=175 Hz) who were also vocalizing digits. Digits (1-9, excluding the bisyllabic ‘se-ven’) were pseudorandomly selected for each speaker such that each speaker produced a distinct digit at any given time. Stimuli were presented diotically through calibrated circumaural headphones (Bose AE2). After familiarization with the task, participants performed randomly interleaved blocks where four blocks had a signal-to-noise ratio of 9 dB SNR, and four blocks at 0 dB SNR. A block consisted of 10 trials (each a 4-digit sequence), where the first two trials were adaptive in difficulty, designed to re-familiarize the participant with the target, and were excluded from analyses. In each trial, digits were spaced with 0.68 s between onsets, and a virtual keypad appeared following the fourth digit to allow participants to report the target digits. Trials where participants took more than 6 seconds to make an initial response were disregarded (on average, less than 1 trial per person). The target speaker was presented at 65 dB SPL. Feedback was not provided during testing. Simultaneously, participants fixated on a cross in the center of a front-facing monitor whilst pupil diameter was recorded. Experimental conditions matched those of the light reflex stimulus, aside from constant isoluminant levels.

### The Speech, Spatial, and Qualities of Hearing Scale

The lived experience of listening in everyday situations was assessed with the Speech, Spatial, and Qualities of Hearing Scale (SSQ; (47)). Forty-nine questions across three sections were scored between 0 and 10 on a 100-interval scale. Responses were provided via a tablet computer (Microsoft Surface Pro, Redmond, WA). Four participants had missing or incomplete SSQ questionnaires.

### Pupillometry data analysis

Blinks were linearly interpolated across and pupil traces were downsampled to 20 Hz. Trials where participants blinked in the second preceding stimulus onset were disregarded. Pre-stimulus pupil diameter was quantified as the mean value between -1 and 0 seconds relative to the onset of the first digit stimulus. For each participant, pre-stimulus pupil diameters were quintiled, i.e., discretized across 5 equally sized rank-ordered bins. Data from individuals who had bins with fewer than 2 values in were excluded (N=18, an additional 4 participants had blink-contaminated values clustered towards the start or end of the task and were excluded from time-related analysis [i.e., Fig. 2B]). We also calculated z-score normalized values with the mean and standard deviation of each participants’ pre-stimulus pupil diameters. Pre-stimulus pupil diameters were also characterized as a proportion of pupil size extremes during a dynamic light stimulus. For this stimulus, we again interpolated across blinks and determined individual pupil size limits by calculating the lowest and highest percentile values observed (due to unsuccessful tracking during the light stimulus, ten participants’ pupil data not unable to undergo this final transformation).

### Statistical analysis

Statistical analyses were performed with MATLAB (Mathworks, Natick, MA). Arcsine transformations were applied to mean digits-in-noise scores prior to statistical testing, to counteract small variance near 0 and 100%. To quantify the dependence of digits-in-noise behavior on pre-stimulus pupil diameter, we calculated mean accuracy and response times for 0 dB SNR trials associated with each pre-stimulus pupil quintile. For this analysis approach we first used a polynomial regression (first order) to control for any influence of trial number on behavioral outcomes. We quantified non-monotonicity as the mean change in behavioral performance between the 3^rd^ and 1^st^/5^th^ quintiles and tested whether population means were non-zero (either two-sided Student’s t-tests or Wilcoxon rank-sum tests, dependent on normality assessments).

We used mediation analysis to test whether PTA (0.5,1, and 2 kHz average) statistically mediated the influence of age on (i) mean multi-talker speech intelligibility, or (ii) changes in speech intelligibility at PPAS extremes. We used the M3 Mediation Toolbox (71, 72) to derive a bootstrapped distribution for the indirect effect (i.e., via PTA), providing 95% confidence intervals to test the *a*x*b* coefficient.

We also fit a statistical model to the behavioral data – a generalized linear mixed-effects (GLME) model. In the GLME, binomial outcomes in each digits-in-noise 0 dB SNR trial were modelled as a linear combination of predictors passed through a probit link function. Age, PTA, PPAS^2^, and trial number were treated as fixed effects, with an interaction term for age and PPAS^2^, and for PTA and PPAS^2^. Study participant was included as a random effects intercept. The model formula was specified as follows: OUTCOME ∼ (PPAS^2^)*(AGE+PTA)+TRIAL+(1|PARTICIPANT).

To determine the influence of age group and another categorical variable (e.g., signal-to-noise ratio) on a continuous outcome measure (e.g., evoked-pupil size) we ran two-way analysis of variance testing.

We fit a multi-variate linear regression model with five predictor variables (for each study participant: pure-tone average, mean accuracy, mean change in accuracy between the 3^rd^ and 1^st^ pre-stimulus pupil quintiles, mean change in accuracy between the 3^rd^ and 5^th^ pre-stimulus pupil quintiles, mean evoked pupil size [% of light range]) and individual SSQ scores as the response variable. The model was fit via a QR decomposition algorithm using the MATLAB function ‘fitlm’.

Dispersion factors were derived from binomial fits to participant-level 0 dB SNR trial-level data using the MATLAB function ‘fitglm’. Dispersion value larger than 1 indicate overdispersion relative to a binomial distribution.

To assess statistical significance, we used a p-value criterion of p<0.05 (symbolized with a single asterisk, p<0.01 symbolized by two asterisks, and p<0.001 symbolized by three asterisks). Specific statistical details can be found in the main text and corresponding figure legends.

## Acknowledgments

This research was supported by a grant from the National Institute on Deafness and Other Communication Disorders [P50DC015857 (D.B.P.)] and a Misophonia Research Foundation Impact Award (S.S.S.). We thank Kenneth Hancock and Kelly Jahn for supporting initial hardware and software development.

