## Supplemental Materials for "Arousal state fluctuations are a source of internal noise underlying age-related declines in speech intelligibility"

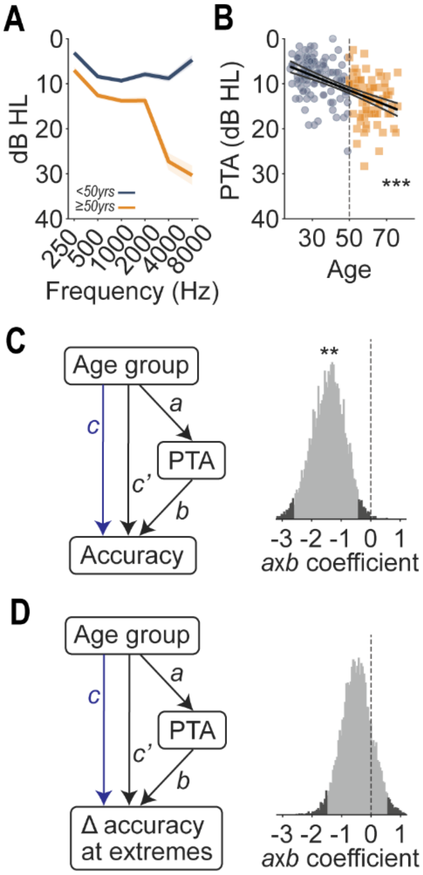


**Figure S1.** Examining pure-tone averages as a potential mediator. (A) Mean hearing thresholds split by the younger and older adults (<50, blue; ≥50, orange). Shaded region denotes standard error. (B) Age was negatively correlated with pure-tone averages (PTA; Pearson's r=0.52, *P*<0.001, n = 183). (C) A mediation analysis found that PTA statistically mediated the influence of age group on mean speech intelligibility performance. *Right,* bootstrapped regression coefficients of the indirect path (bootstrapped p-value for the indirect effect *axb* was *P*=0.004). Dark/light shading of the histogram shows 95% confidence interval for *axb.* (D) As per (C), but by contrast to the influence of PTA on overall speech accuracy, PTA did not mediate the effect of age on accuracy changes at pupil-indexed extremes (bootstrapped p-value for the indirect effect *axb* was *P*=0.39).

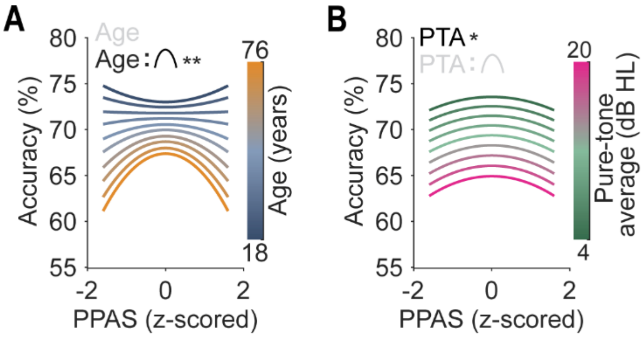


**Figure S2.** A regression analysis emphasizes the interaction of PPAS with age**.** Performance estimates from a generalized linear mixed-effects model. The two panels depict the effects of (A) age and (B) pure-tone average, and their relationship with PPAS, whilst other factors are held constant. The only factor that had a significant interaction with PPAS was age, which produced an increasingly negative quadratic function at older ages. Significant model terms are shown in black.

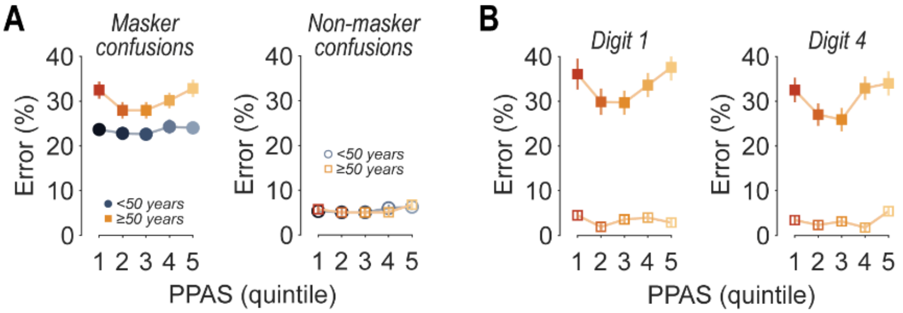


**Figure S3.** Errors are largely due to reporting competing speech**.** (A) Dependence of errors on PPAS, where a digit spoken by a competing talker is misattributed as the target (left, “masker confusion”) or participants reported an unspoken digit (right, “non-masker confusion”. Error likelihoods are shown averaged for both younger (blue) and older (orange) adults, where error bars denote standard error. (B) The relationship between masker confusions and pupil size showed non-monotonic trends for the initial digits and for the final digit in older adults.

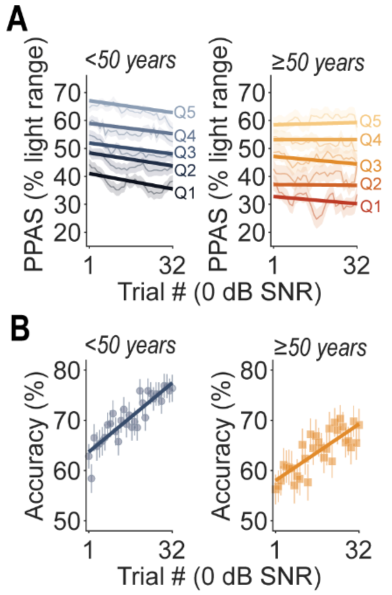


**Figure S4.** Changes in PPAS and performance by trial. (A) We calculated PPAS quintiles benchmarked against light responses, with an 8-trial sliding window. Overlaid are median linear regression fits. Results echo Fig. 2A, with declining PPAS values with trial number for younger adults (blue, left panel), but not for the cohort of older adults (orange right panel). (B) Task performance improved over the 32 trials at 0 dB SNR for both the younger (blue, left panel) and older adults (orange, right panel).

| Predictor | FStat (DF1, DF2) | p-value |
|---|---|---|
| Age | 1.56 (1, 4352) | 0.212 |
| PTA | 4.82 (1, 4352) | **0.028** |
| Trial | 105.7 (1, 4352) | **<0.001** |
| $PPAS^2$ | 4.47 (1, 4352) | **0.035** |
| $PPAS^2$ x Age | 6.90 (1, 4352) | **0.009** |
| $PPAS^2$ x PTA | 0.034 (1, 4352) | 0.852 |

**Table S1.** Generalized linear mixed-effects model ANOVA.
